# A Fob1 independent role for the Smc5/6 complex in the rDNA that modulates lifespan

**DOI:** 10.1101/2020.01.24.917203

**Authors:** Sarah Moradi-Fard, Aditya Mojumdar, Megan Chan, Troy A. A. Harkness, Jennifer A. Cobb

## Abstract

The ribosomal DNA (rDNA) in *Saccharomyces cerevisiae* is in one tandem repeat array on Chromosome XII. Two spacer regions within each repetitive element, called <u>n</u>on-transcribed <u>s</u>pacer <u>1</u> (NTS1) and NTS2, are important in nucleolar organization. Smc5/6 localizes to both NTS1 and NTS2 and is involved in the regulation of Sir2 and Cohibin binding at NTS1, whereas Fob1 and Sir2 are required for optimal binding of the complex to NTS1 and NTS2, respectively. We demonstrate that Smc5/6 functions in chromatin silencing at NTS1 independently of its role in homologous recombination (HR) when forks pause at the replication fork barrier (RFB). In contrast, when the complex does not localize to the rDNA in *nse3*-1 mutants, the shortened lifespan correlates with NTS2 homeostasis independently of *FOB1* status. Our data identify the importance of Smc5/6 integrity in NTS2 transcriptional silencing and repeat tethering, which in turn underscores rDNA stability and replicative lifespan.

**Highlights:**

- Smc5/6 is important for transcriptional silencing in the rDNA.
- Smc5/6 tethers the rDNA array to the periphery.
- Transcriptional silencing of ncRNA at NTS1 and NTS2 is differentially regulated.
- Smc5/6 has a role in rDNA maintenance independent of HR processing at the RFB.
- Fob1-independent disruption of Smc5/6 at NTS2 leads to lifespan reduction.

## INTRODUCTION

The ribosomal DNA in *S. cerevisiae* (budding yeast) consists of approximately 150-200 identical 9.1 kb long tandem repeats on chromosome XII which are assembled in one cluster and positioned close to the nuclear periphery^1^. The 35S and 5S ribosomal RNA genes are transcribed by RNA polymerases I and III respectively and two spacer sequences flanking the rRNA genes can be transcribed by RNA polymerase II to produce noncoding (nc) RNAs^2^. The intergenic regions are usually silenced and referred to as <u>n</u>on-transcribed <u>s</u>pacer <u>1</u> (NTS1) and NTS2 (Fig. 1a)^3, 4^. NTS1 contains a replication fork barrier (RFB) sequence and a bi-directional non-coding promoter, called E-pro, and NTS2 contains an autonomous replication sequence (ARS). The histone deacetylase Sir2 interacts with Net1 and Cdc14 in the nucleolus to form the RENT complex, which represses transcription from NTS1 and NTS2^3–13^. The recovery of Sir2 at NTS2 appears to be dynamic and dependent on RNA Polymerase I transcription of 35S, which is found at ∼50% of rDNA genes in asynchronously growing cell cultures^14, 15^. The recovery of Sir2 at NTS1 is through another mechanism that has been characterized more extensively compared to its binding at NTS2. At NTS1, the Fob1 protein binds the RFB, ensuring unidirectional replication and the localization of RENT, which represses E-pro transcription. When the progressing replication fork is stalled by Fob1, a double strand break (DSB) results and it is repaired by recombination between the repetitive sequences^16–18^. From an evolutionary perspective, the events at NTS1 are important to maintain rDNA copy number as DSB repair can occur through unequal sister chromatid recombination (USCR), allowing changes in the number of repetitive elements (contraction or expansion)^19^. Increased transcription from the E-pro, loosens chromatin adjacent to the DSB induced at the RFB, which in turn leads to increased USCR-mediated repair. However, if this process is not highly regulated then rDNA instability ensues. Indeed, the loss of *SIR2* results in transcription from NTS1 and rDNA instability^20, 21^. Importantly, many *sir2*Δ mutant phenotypes in the rDNA depend on the formation of a DSB at the RFB and are reversed through the additional deletion of *FOB1*.

**Fig. 1.**
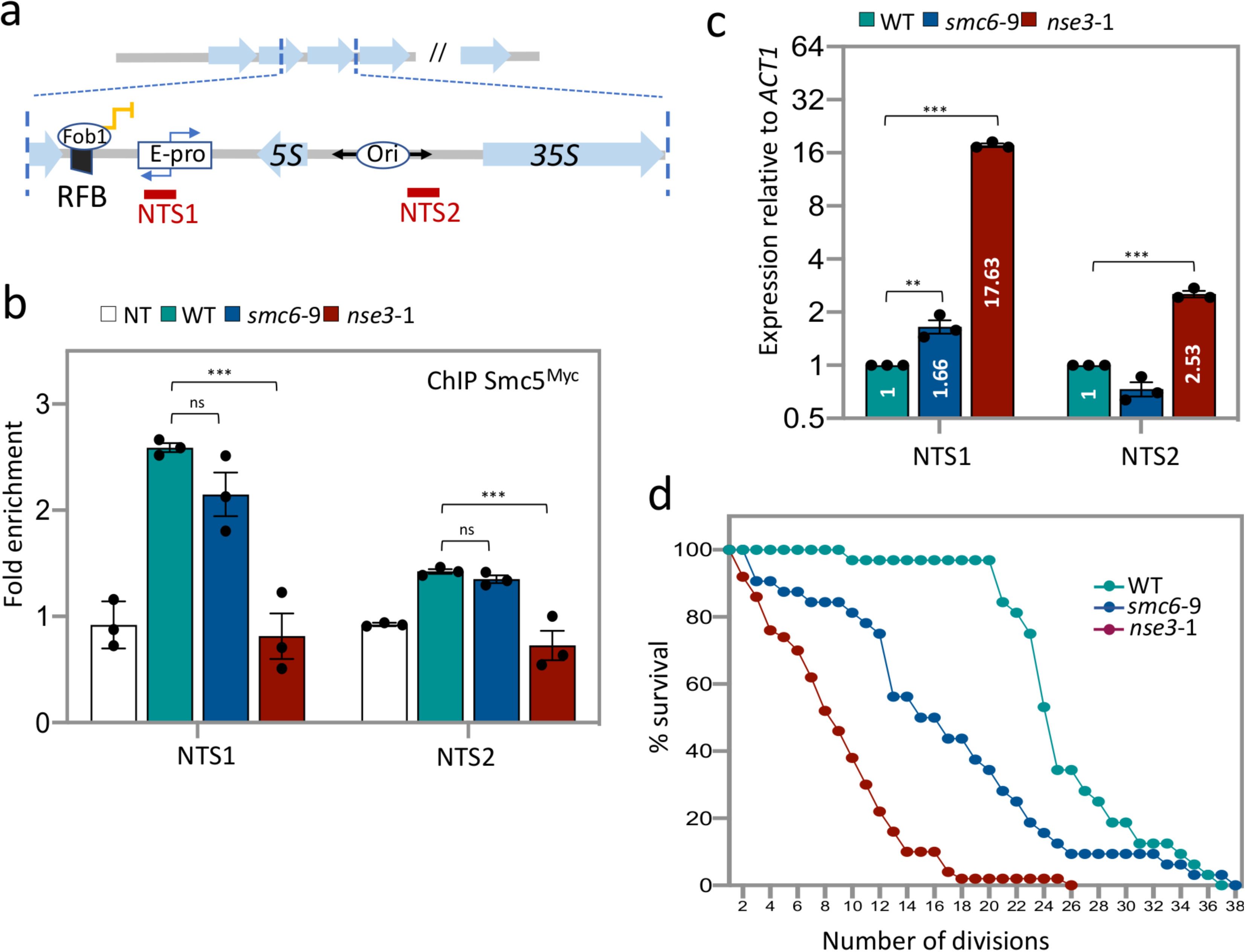
Smc5/6 localization to NTS1 and NTS2 is important for lifespan. **a** Schematic of rDNA repeats in *S. cerevisiae* showing non-transcribed spacers (NTS1 and NTS2) flanking the transcribed 5S and 35S sequences in one repeat. The location of primer sites used in ChIP experiments are illustrated. **b** Enrichment of Smc5^Myc^ at NTS1 and NTS2 by ChIP with α-Myc in non-tagged control (JC 470), WT (JC 3467), *smc6*-9 (JC 5039) and *nse3*-1 (JC 3483) at NTS1 and NTS2. Fold enrichment is based on normalization to negative control region (ZN) as previously described^44^. **c** Transcription at NTS1 and NTS2 relative to WT cells after normalization to *ACT1* transcription for WT (JC 471), *smc6*-9 (JC 1358) and *nse3*-1 (JC 3032). **d** Replicative lifespan measured and represented as percentage of survival of mother cells with each division for WT (JC 471), *smc6*-9 (JC 1358) and *nse3*-1 (JC 3032) strains. Analysis was performed using at least three biological replicates. Statistical analysis is described in methods.

In addition to the RENT complex, a cohesin related complex called Cohibin, consisting of Csm1 and Lrs4, is involved in NTS1-specific silencing independently of Sir2^7, 22–25^. Cohibin physically associates with inner nuclear membrane (INM) proteins Heh1 and Nur1, which are members of the CLIP (chromosome linkage INM proteins) complex^7, 24^. Loss of Cohibin or CLIP components results in defective silencing and tethering of repetitive elements to the periphery^24^. However, loss of Heh1 does not affect silencing of rDNA and only abrogates rDNA tethering which makes rDNA repeats vulnerable to homologous recombination (HR) by exposing them to HR factors present in the nucleoplasm^24^. Thus, both silencing and tethering are essential for rDNA repeat stability^7, 24^.

In all, the rate of rDNA recombination is tightly regulated by multiple mechanisms including chromatin condensation, transcriptional silencing and spatial organization by anchoring the repetitive elements to the inner nuclear membrane^26^. Related to this, is the observation that decreased rDNA stability is linked to a decreased lifespan^27–29^, which is the number of times a mother cell can bud and give rise to daughter cells before it dies^30–32^. Cells lacking *SIR2* show a decrease in lifespan. Initial correlations indicated that the increased recombination in *sir2*Δ cells, which resulted in an increase rDNA repeats-containing extra chromosomal circles (ERCs), caused premature senescence in the mother cells by titrating limited replication and transcription factors from the genome^29^. Subsequent work however supports a model whereby rDNA instability itself drives aging and that ERC accumulation is not the cause of a decreased lifespan, but rather an indicator of rDNA stability^33^. Importantly, while rDNA stability is a central factor in longevity, the reduced lifespan of *sir2*Δ mutants is not completely suppressed by deleting *FOB1*, which indicates that Sir2 contributes to rDNA stability beyond its known function(s) at NTS1.

The Smc5/6 complex belongs to the SMC family and it is constitutively present at rDNA repeats^34–36^. All components of SMC complexes are essential for life, therefore investigating their functions *in vivo* has relied heavily on characterizing thermosensitive (ts) mutants, which in most cases limits understanding to only a subset of functions. Most of the work with Smc5/6 at the rDNA has linked the complex to HR-resolution of replication forks at the RFB^35–38^. Specifically, in *smc6*-9 mutants, cells display delayed rDNA replication, increased chromosomal breakage and accumulated X-shaped DNA structures^35, 38^. Replication and HR-related defects have also been reported using degron-inducible mutants^37^. The accumulation of HR intermediates in ts and degron-tagged Smc5/6 complex mutants was reversed by deleting *FOB1*^37, 38^. These observations, together with other HR-related functions reported with Smc5/6^39–41^, indicate that Smc5/6 is integral for controlling HR-mediated processes at the rDNA. In line with the described functionalities of the other SMC complexes, cohesion and condensin, at multiple genomic loci^42–44^, HR-independent roles for Smc5/6 in the higher-order organization of repetitive elements has been observed at telomeres ^44^. However, involvement of the Smc5/6 complex in rDNA repeat stability independent of HR has not been reported.

We characterize the loss of Smc5/6 localization in the rDNA utilizing *nse3*-1 mutant cells, which was previously shown to disrupt complex recruitment to telomeres. TPE and clustering were defective in *nse3*-1, but not *smc6*-9 mutants, which were HR deficient but telomere associated^44^. In *nse3*-1 cells, the binding of Smc5/6 to the rDNA array is not detected and loss of the complex leads to multiple rDNA phenotypes, such as increased nucleolar volume, defective silencing, release of rDNA from nuclear periphery and a reduced lifespan. In addition, we show that the Smc5/6 complex is partially required for binding of Cohibin and Sir2 specifically at NTS1, while Fob1 and Sir2 were required for optimal binding of the Smc5/6 to NTS1 and NTS2, respectively. Importantly, shortened lifespan of *smc6*-9 cells (HR-defective mutant) but not *nse3*-1 cells were rescued by the deletion of *FOB1*. The reduced replicative lifespan of *nse3*-1 *fob1*Δ double mutants correlates with loss of Smc5/6 recovery at NTS2, increased transcription at NTS2, persistence of ERC molecules and enlarged nucleolar morphology. In all, our data reveal role(s) for the Smc5/6 complex in nucleolar organization, the proper formation of canonical silencing factors at rDNA as well as interactions with the nuclear periphery independent of fork pausing at the RFB. Particularly, our data strongly suggest the importance of NTS2 events in rDNA stability and replicative lifespan.

## RESULTS

### Absence of the Smc5/6 complex at rDNA results in silencing defects and short lifespan

The Smc5/6 complex binds NTS1 and NTS2 in the rDNA. Previous work with a ts allele of *SMC6*, *smc6*-9, showed that the complex is important for processing HR intermediates that arise when replication forks stall at RFBs in NTS1^35, 38^. The stability of rDNA correlates with lifespan and depends on transcriptional silencing; however the impact of Smc5/6 in these processes has not been reported. Here we characterize *smc6*-9 and another ts allele, *nse3*-1, which disrupts telomere clustering at the nuclear periphery and fails to localize Smc5/6 to telomeres^44^. To determine complex binding, we performed chromatin immuno-precipitation (ChIP) with Smc5^Myc^ followed by qPCR with primers designed to NTS1 and NTS2 (Fig. 1a). Similar to previous reports, Smc5^Myc^ was enriched in the rDNA at both NTS sites (Fig. 1b)^35^. The level of Smc5^Myc^ in *smc6*-9 was similar to wild type indicating that the rDNA defects previously reported with this allele do not stem from defects in the complex binding in spacers regions. In contrast, at NTS1 and NTS2, there was a significant reduction of Smc5^Myc^ in *nse3*-1 mutant cells, to levels indistinguishable from the non-tagged control (Fig. 1b). Not only was Smc5^Myc^ reduced at NTS regions, but also at sites in the 35S and 5S ribosomal RNA genes in *nse3*-1 mutant cells, indicating that *nse3*-1 potentially imposes a global DNA-binding defect for the Smc5/6 complex in the rDNA (Supplementary Fig. 1b). Proper regulation of chromatin silencing in the NTS sites (mostly studied for NTS1) correlates strongly with rDNA stability and lifespan^3–13, 21–25^, thus we concentrated on potential functions for Smc5/6 together with known factors involved in rDNA stability. Therefore, we reasoned that further comparisons between *nse3*-1 and *smc6*-9 alleles could reveal functions for the Smc5/6 complex that could distinguish its role in HR processing at the RFB from other potential functions in the rDNA. As such, the levels of ncRNAs at NTS1 and NTS2 were measured in the mutant alleles as previous work showed that silencing at sub-telomeric loci was defective when the complex was not recovered at telomeres in *nse3-*1 mutant cells^44^. There was ∼ 2-fold increase in expression at NTS1 in *smc6*-9 mutants compared to wild type (Fig. 1c). Transcription in *nse3-*1 mutants was markedly higher at both sites, indicating a role for Smc5/6 in NTS2 repression which is regulated differently than at NTS1 (Fig. 1c). Repression of ncRNA from the E-pro located in NTS1 has been linked to lifespan as increased transcription correlates with decreased lifespan^21^. Compared to wild type, cells harboring either mutant alleles showed a shorter lifespan, however it was reduced more in *nse3*-1 cells, when the complex is unable to bind chromatin (Fig. 1d).

### Smc5/6 complex interacts with CLIP and is required for Heh1-mediated rDNA tethering and Heh1-independent rDNA compactness

Previous work in budding yeast showed that an increase in replicative aging correlated with an increase in nucleolar volume^45^. Therefore, we visualized the morphology of the nucleolus in wild type and mutant cells with Nop1^CFP^ marking the nucleolus and Nup49^GFP^ marking the nuclear periphery (Fig. 2a). In *smc6*-9 mutant cells, nucleolar volume slightly increased, ∼ 1.14-fold, whereas in *nse3*-1 mutant cells the volume of the nucleolus was almost twice as large as in wild type cells (Fig. 2b). These data correlate Smc5/6 binding at the rDNA with maintaining nucleolar compaction.

**Fig. 2.**
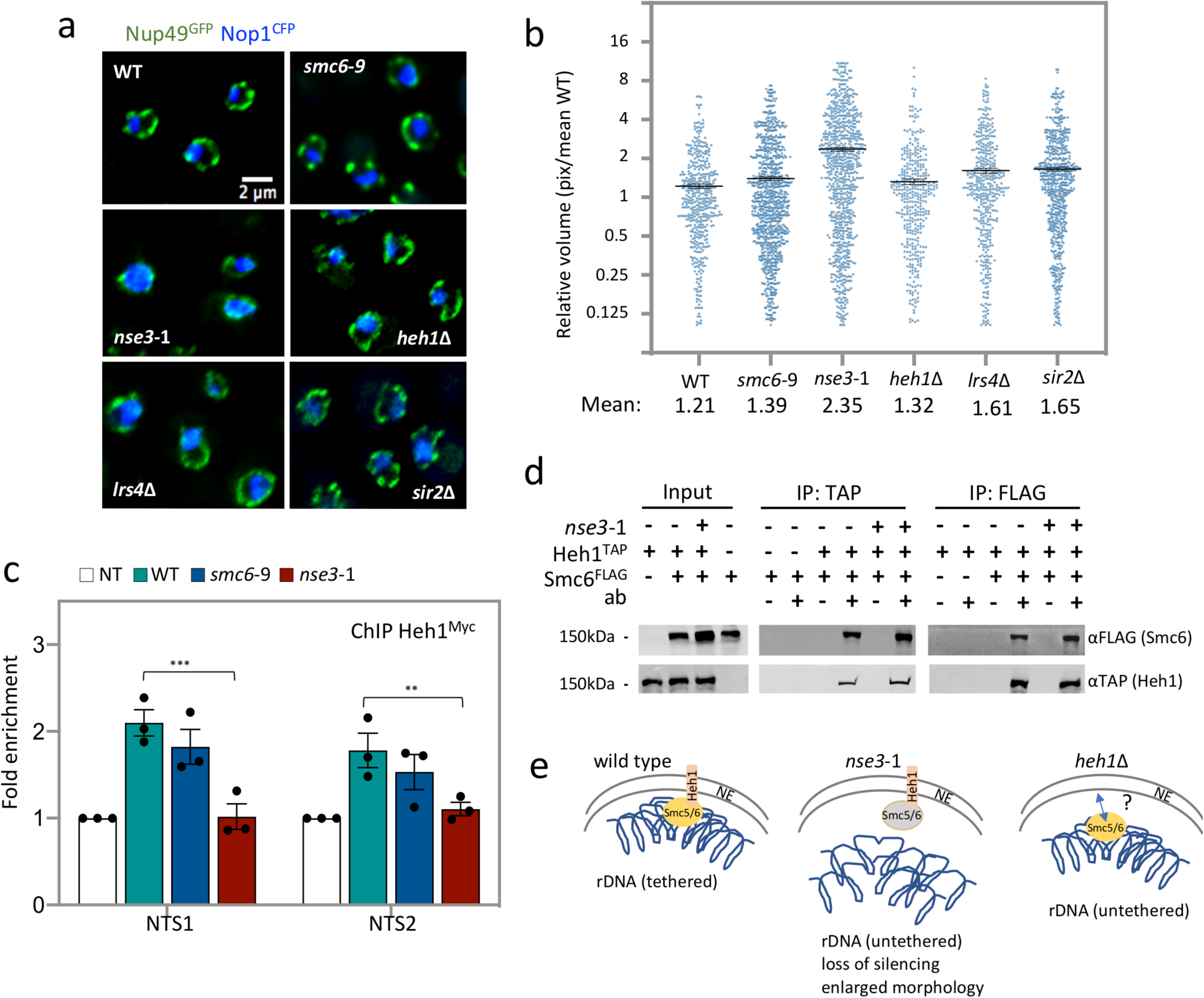
Smc5/6 tethers rDNA repeats at the periphery and interacts with Heh1. **a** Nucleolus morphology is illustrated by imaging CFP-tagged NOP1 in WT (JC 4676), *smc6*-9 (JC 4932), *nse3*-1 (JC 4729), *heh1*Δ (JC 4735), *lrs4*Δ (JC 4731) and *sir2*Δ (JC 4633); GFP-tagged NUP49 indicates nuclear periphery boundaries. **b** Scatter plot data of nucleolar volume for WT (JC 5016), *smc6*-9 (JC 5014), *nse3*-1 (JC 5015), *heh1*Δ (JC 4735), *lrs4*Δ (JC 4731) and *sir2*Δ (JC 4633) were measured in pixel and represented relative to mean of WT as described in methods. **c** Enrichment of Heh1^Myc^ at NTS1 and NTS2 by ChIP with α-Myc in non-tagged control (JC 470), WT (JC 4022), *nse3*-1 (JC 4228) and *smc6*-9 (JC 4942) at NTS1 and NTS2. Fold enrichment is represented as relative to no tag control after normalization to negative control region (ZN). **d** Co-IP between Smc6-FLAG and Heh1-TAP followed with western blotting using corresponding antibodies to epitope tags on each protein. Heh1-TAP IP and Smc6-FLAG IP was performed in negative control (JC 1594), WT (JC 4811) and *nse3*-1 (JC 4813). **e** Schematic representation of Smc5/6 in rDNA tethering at the periphery in i. wild type, ii. *nse3*-1, and iii. *heh1*Δ cells. Analysis was performed using at least three biological replicates. Statistical analysis is described in methods.

Given the altered morphology of the nucleolus in *nse3*-1 mutants and previous work showing Smc5/6 localizes at nuclear periphery^46^, we examined a potential role of the complex in anchoring the rDNA to the inner nuclear membrane (INM). Heh1 and Nur1 reside in the INM and form the CLIP complex. The recovery of Heh1 in the rDNA by ChIP has been used to measure anchoring of the repeats at the nuclear periphery^24^. Compared to wild type, Heh1^Myc^ enrichment at NTS1 and NTS2 decreased significantly in *nse3*-1, but not *smc6*-9 mutants (Fig. 2c). Co-immunoprecipitations (IPs) between Heh1^TAP^ and Smc6^FLAG^ showed an interaction between Smc5/6 and Heh1 under physiological condition that was not diminished in cells harboring the *nse3*-1 allele (Fig. 2d). Deletion of *HEH1* did not alter Smc5/6 enrichment in the rDNA (Supplementary Fig. 2) and nucleolar compaction in *heh1*Δ mutants was similar to wild type (Fig. 2a, b). Thus, Smc5/6 is important for both rDNA compaction and for tethering the rDNA repeats to the periphery through interactions with Heh1 in the CLIP complex (Fig. 2e), however our data suggest that Heh1-dependent tethering and nucleolar compaction are separable (Fig. 2e).

### Smc5/6 physically and genetically interacts with Sir2 and Cohibin

Cohibin and the RENT complex also interact with CLIP and contribute to rDNA repeat tethering^7, 24^. The two components of the Cohibin complex, Lrs4 and Csm1, interact with Sir2 as part of the RENT complex and both are present at the rDNA and the nuclear periphery^7, 23–25, 47^. In a side-by-side comparison the deletion of either *LRS4* or *SIR2* led to increased nucleolar morphology, however the increase was below that measured in *nse3*-1 mutant cells (Fig. 2a, b). In addition to the morphological changes, transcriptional silencing is another pathway where Smc5/6, Cohibin and Sir2 function (Fig. 1c)^24, 47^, thus we wanted to investigate the interplay between Smc5/6 and these the canonical silencing factors in rDNA homeostasis^3–13, 22–25^. ChIP was performed with Csm1^TAP^ to measure Cohibin recovery in the rDNA. In *nse3*-1 cells there was a 3-fold reduction in Csm1^TAP^ enrichment at NTS1 (Fig. 3a). Cohibin binding in NTS2 was reduced overall compared to NTS1, and this was further decreased in *nse3*-1 mutants, whereas Csm1^TAP^ recovery in *smc6*-9 at either NTS region was indistinguishable from wild type (Fig. 3a). In all, the recovery of Cohibin at the rDNA was partially dependent on Smc5/6. To determine whether a physical interaction could be detected *in vivo* between Smc5/6 and Cohibin, co-IP was performed between the Csm1^TAP^ and Smc6^FLAG^. Smc6^FLAG^ was recovered in α-TAP (Csm1) pulldowns and vice versa, Csm1^TAP^ was recovered in α-FLAG (Smc6) IPs (Fig. 3b). Similar co-IP experiments in *nse3*-1 mutant cells showed unchanged interactions between the Smc5/6 and Cohibin (Fig. 3b). Yeast two hybrid (Y2H) experiments between the Cohibin subunits, Lrs4 and Csm1, and multiple components of the Smc5/6 complex showed physical interactions between Lrs4 and Nse6 and between Csm1 and both Nse6 and Mms21 (Fig. 3c, Supplementary Fig. 3). Interestingly, Mms21 interacted with Heh1 by Y2H too (Fig. 3c, Supplementary Fig. 3), which prompted us to also determine whether the interaction between Cohibin and the CLIP complex might be mediated by Smc5/6 at the rDNA. Consistent with previous reports, by co-IP we observed interactions between Csm1^TAP^ and Heh^Myc^ (Fig. 3d)^7, 24^. Binding was not altered in *nse3*-1 mutant cells (Fig. 3d), indicating that like Smc5/6 (Fig. 2d), Cohibin interacts with Heh1 independently of its localization at the rDNA array.

**Fig. 3.**
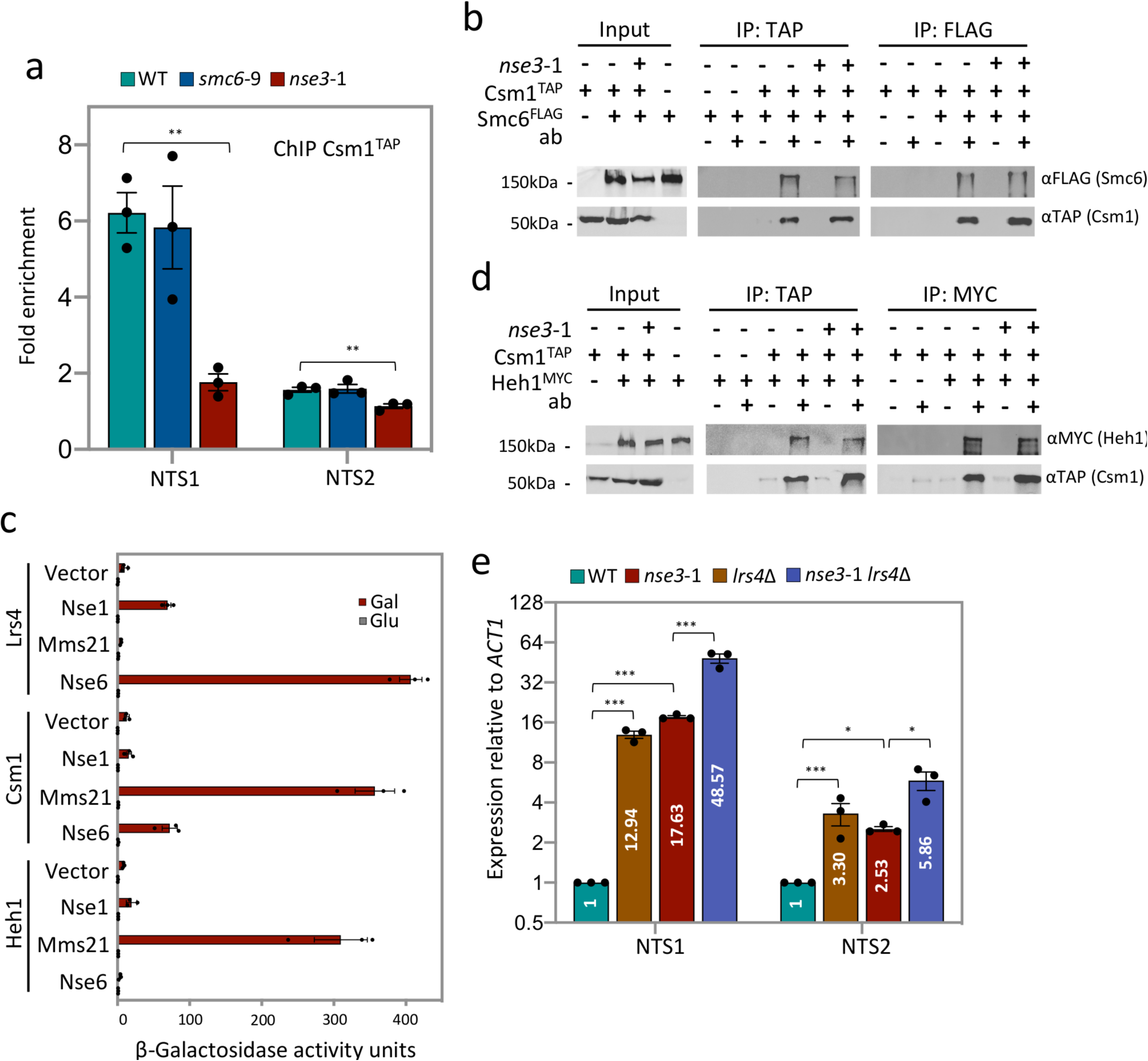
Cohibin recovery at the rDNA is partially dependent on Smc5/6. **a** Enrichment of Csm1^TAP^ at NTS1 and NTS2 by ChIP with α-TAP in WT (JC 4233), *smc6*-9 (JC 4938) and *nse3*-1 (JC 4251) at NTS1 and NTS2. Fold enrichment is based on normalization to negative control region (ZN). **b** Co-IP between Smc6-FLAG and Csm1-TAP followed with western blotting using corresponding antibodies to epitope tags on each protein. Smc6-FLAG IP and Csm1-TAP IP was performed in negative control (JC 1594), WT (JC 4598) and *nse3*-1 (JC 4712). **c** Yeast-two Hybrid analysis between Smc5/6 (Nse6 Nse3 and Mms21), Cohibin (Lrs4 and Csm1) components and Heh1 using quantitative β-galactosidase activity assay. **d** Co-IP between Csm1-TAP and Heh1-^Myc^ followed with western blotting using corresponding antibodies to epitope tags on each protein. Csm1-TAP IP and Heh1-^Myc^ IP was performed in negative control (JC 4224), WT (JC 4774) and *nse3*-1 (JC 4773). **e** Transcription at NTS1 and NTS2 relative to WT cells after normalization to *ACT1* expression for WT (JC 471), *lrs4*Δ (JC 3791), *nse3*-1 (JC 3032) and *nse3*-1 *lrs4*Δ (JC 3796). Analysis was performed using at least three biological replicates. Statistical analysis is described in methods.

As both Smc5/6 and Cohibin contribute to silencing (Fig. 1c)^7^, levels of transcription were compared at the NTS regions in *nse3*-1 and *lrs4*Δ single and double mutant cells. The loss of silencing was additive in *nse3*-1 *lrs4*Δ double mutant cells compared to the single mutant counterparts (Fig. 3e), suggesting one or both of these complexes could impact silencing through another mechanism. Sir2 as part of the RENT complex is important in rDNA transcriptional silencing and its deletion was assessed together with *nse3*-1 and *lrs4*Δ. The impact of the various double and triple mutant combinations was different at NTS1 and NTS2. At NTS1, transcription was ∼2-fold higher in double mutants where *SIR2* was deleted, *nse3*-1 *sir2*Δ and *lrs4*Δ *sir2*Δ, compared to *nse3*-1 *lrs4*Δ mutants (Fig. 4a). Transcription in triple mutant cells was even further increased, where the loss of silencing was markedly greater than in any double mutant combination (Fig. 4a). At NTS2, transcription in *sir2*Δ was ∼ 4-fold higher than in *nse3*-1 or *lrs4*Δ mutant cells, but there was no additive effect in double mutants containing the *SIR2* deletion compared to sir2Δ single mutants (Fig. 4a). NTS2 transcription in *nse3*-1 *lrs4*Δ *sir2*Δ triple mutant cells increased ∼2-fold above *nse3*-1 *sir2*Δ and *lrs4*Δ *sir2*Δ double mutants and ∼8-fold above *nse3*-1 *lrs4*Δ, but loss of silencing at NTS2 in the triple mutant cells was not synergistic like at NTS1 (Fig. 4a).

**Fig. 4.**
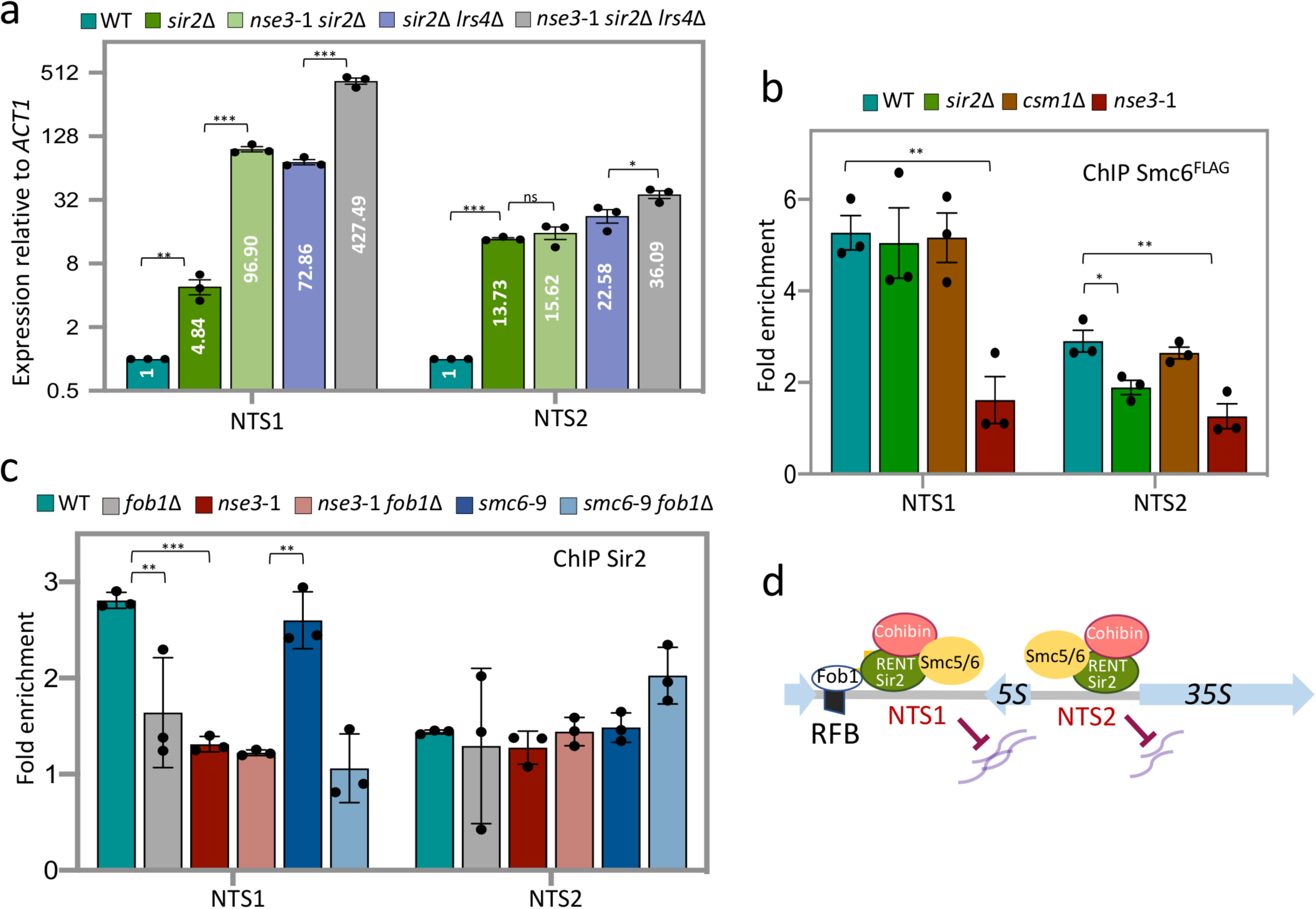
Smc5/6 together with Sir and Cohibin silence NTS1 and NTS2. **a** Transcription of NTS1 and NTS2 relative to WT cells after normalization to *ACT1* expression for WT (JC 471), *sir2*Δ (JC 4648), *nse3*-1 *sir2*Δ (JC 3787), *sir2*Δ *lrs4*Δ (JC 4979) and *nse3*-1 *sir2*Δ *lrs4*Δ (JC 4980). **b** Enrichment of Smc6^FLAG^ at NTS1 and NTS2 by ChIP with α-FLAG in WT (JC 1595), *sir2*Δ (JC 4699), *csm1*Δ (JC 4243) and *nse3*-1 (JC 3078). Fold enrichment is based on normalization to negative control region (ZN). **c** Enrichment of Sir2 at NTS1 and NTS2 by ChIP with α-Sir2 in WT (JC 471), *fob1*Δ (4825), *nse3*-1(JC 3032), *nse3*-1 *fob1*Δ (JC 4595), *smc6*-9 (JC 1358) and *smc6*-9 *fob1*Δ (JC 4824) strains at NTS1 and NTS2. Fold enrichment is based on normalization to negative control region (ZN). **d** Schematic representation of Smc5/6, Sir2 and Cohibin localization and their relationship to transcription at NTS1 and NTS2 in WT vs. *nse3*-1 *sir2*Δ *lrs4*Δ cells. Analysis was performed using at least three biological replicates. Statistical analysis is described in methods.

To correlate transcriptional silencing with the physical presence of these complexes in the rDNA, ChIP was performed on Smc6^FLAG^. Similar to Smc5^Myc^ (Fig. 1b), Smc6^FLAG^ recovery at NTS1/2 was reduced in *nse3*-1 mutant cells (Fig. 4b). In contrast, Smc6^FLAG^ recovery in *csm1*Δ or *sir2*Δ was similar to wild type indicating Smc5/6 recovery at NTS1 is independent of the Cohibin and RENT complexes. Like with Csm1^TAP^ (Fig. 3a), the recovery of Sir2 was reduced at NTS1 in *nse3*-1 mutant cells (Fig. 4c). Taken together, Smc5/6 localization at the rDNA is important for Cohibin and Sir2 association but its contribution to silencing is partially independent as triple mutant cells show increased NTS1 transcription. Smc6^FLAG^ recovery decreased at NTS2 in *sir2*Δ, but not *csm1*Δ mutants (Fig. 4b), indicating that Smc5/6 association with NTS2 is partially dependent on the RENT complex but not Cohibin. This correlates with the epistatic relationship for loss of silencing in *sir2*Δ and *nse3*-1 *sir2*Δ mutants (Fig. 4a). Notably, and in contrast to *nse3*-1, the binding of Csm1^TAP^ and Sir2 were unchanged in *smc6*-9 mutants indicating that the physical presence of Smc5/6, not its HR-linked functions, is critical for recruiting these canonical rDNA silencing factors (Fig. 3a, 4c). Taken together, our data support a model whereby Smc5/6 is physically present at both NTS regions (Fig. 4d), but the interplay of Smc5/6 with Cohibin and RENT at the two regions is regulated differently. For example, the deletion of *SIR2* impacts Smc5/6 at NTS2 but not at NTS1 and the presence of Smc5/6, independently of its function in HR, impacts Sir2 and Cohibin levels more at NTS1.

### Smc5/6 complex plays a Fob1-independent role in modulating lifespan

Both Fob1 and Smc5/6 physically and genetically interact with Sir2^14, 35–37, 48–51^. Sir2 recovery at NTS1 in *smc6*-9 was similar to wild type, however consistent with previous reports^14^ its association at NTS1 was Fob1-dependent as Sir2 was equally reduced in *fob1*Δ and *smc6-*9 *fob1*Δ mutants (Fig. 4c). Expression of ncRNA at NTS1 in *smc6*-9 *fob1*Δ increased relative to either single mutant and was similar to the levels in *sir2*Δ (Fig. 4a, 5a). In contrast, deletion of *FOB1* did not impact the silencing defects in *nse3*-1 mutants at NTS1, nor the level of transcription at NTS2 in either *smc6*-9 or *nse3*-1 (Fig. 5a). Increased nucleolar volume was observed in mutant cells with increased transcription (Fig. 2a, b), therefore we measured the nucleolar volume of *smc6*-9 and *nse3*-1 in combination with *fob1*Δ. *FOB1* deletion in *smc6*-9 mutants resulted in a more compact nucleolus (Fig. 5b). However, *fob1*Δ did not reverse the enlarged nucleolus in *nse3*-1 mutants, but rather nucleolar volume increased in *nse3*-1 *fob1*Δ double mutants (Fig. 5b).

**Fig. 5.**
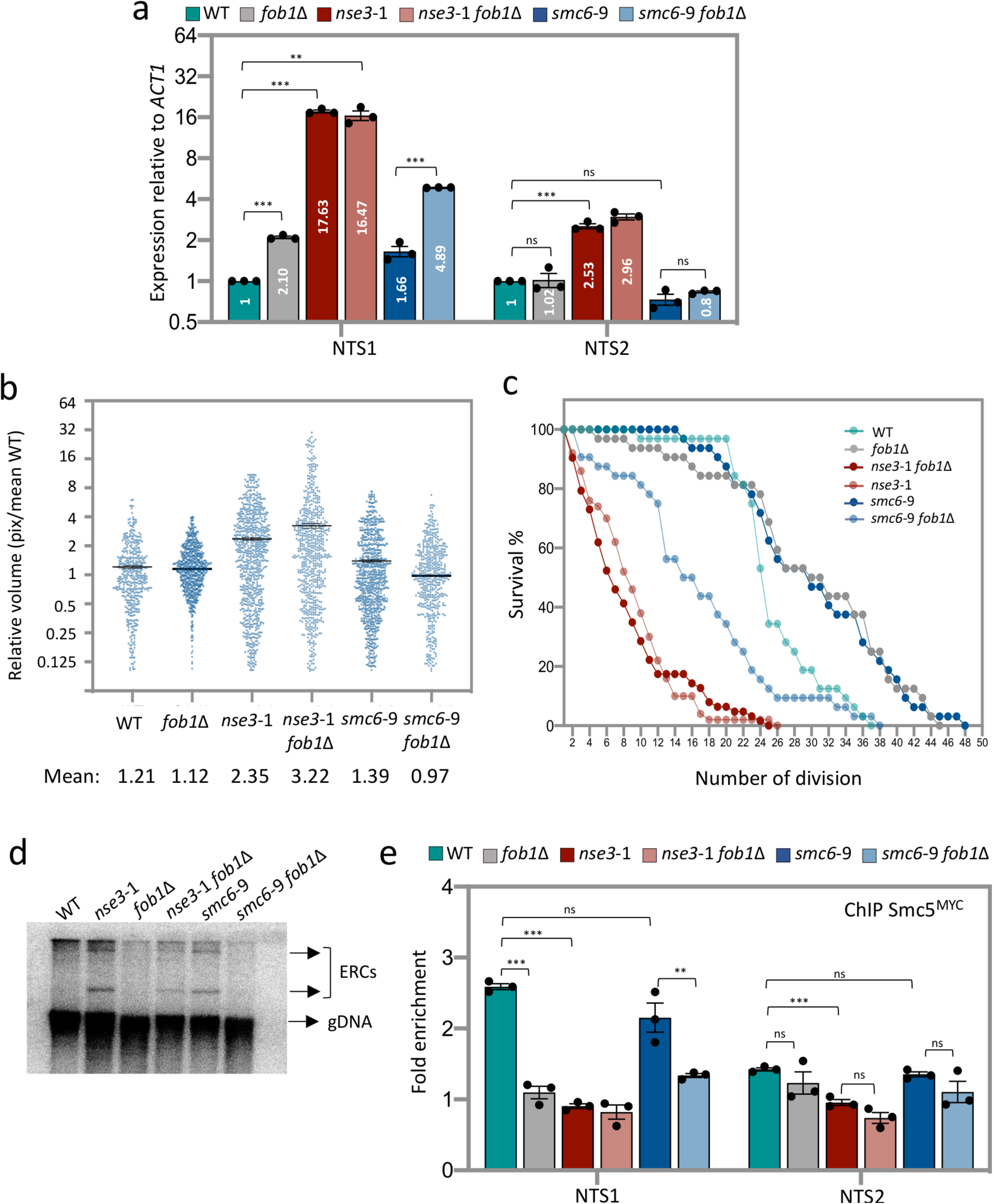
Smc5/6 function at NTS2 is important for nucleolar homeostasis independent of HR processing at the RFB. **a** Transcription of NTS1 and NTS2 measured and represented as relative to WT cells after normalization to *ACT1* expression for WT (JC 471), *fob1*Δ (JC 4825), *nse3*-1 (JC 3032), *nse3*-1 *fob1*Δ (JC 4595), *smc6*-9 (JC 1358) and *smc6*-9 *fob1*Δ (JC 4824) strains. **b** Scatter plot data of nucleolar volume for WT (JC 5016), *fob1*Δ (JC 4985), *nse3*-1 (JC 5015), *nse3*-1 *fob1*Δ (JC 5110), *smc6*-9 (JC 5014) and *smc6*-9 *fob1*Δ (JC 5113) strains were measured in pixel and represented relative to mean of WT. **c** Replicative lifespan measured and represented as percentage of survival of mother cells with each division for WT (JC 471), *fob1*Δ (4825), *nse3*-1 (JC 3032), *nse3*-1 *fob1*Δ (JC 4595), *smc6*-9 (JC 1358) and *smc6*-9 *fob1*Δ (JC 4824) strains. **d** ERC molecules abundance in WT (JC 471), *fob1*Δ (JC 4825), *nse3*-1 (JC 3032), *nse3*-1 *fob1*Δ (JC 4595), *smc6*-9 (JC 1358) and *smc6*-9 *fob1*Δ (JC 4824) strains. **e** Enrichment of Smc5^Myc^ at NTS1 and NTS2 by ChIP with α-^Myc^ in WT (JC 3467), *fob1*Δ (JC 5041); *nse3*-1 (JC 3483), *nse3*-1 *fob1*Δ (JC 5044), *smc6*-9 (JC 5039) and *smc6*-9 *fob1*Δ (JC 5040). Fold enrichment is based on normalization to negative control region (ZN). Analysis was performed using at least three biological replicates. Statistical analysis is described in methods.

Both the *smc6*-9 and *nse3*-1 alleles have a shorter lifespan compared to wild type (Fig. 1d). A shorter lifespan has been attributed to rDNA instability arising from increased collisions between replication and transcription machineries when chromatin was not silenced at NTS1^52^. Fob1 binding at the RFB is central in this process and Smc5/6 is also linked, as it modulates HR processing at stalled replication forks^14, 16, 19, 37, 38, 49, 53, 54^ and here we show transcriptional silencing at NTS1. Previous work has shown that the reduced lifespan of *sir2*Δ mutants is suppressed by deleting *FOB1*^51^. To understand the impact of Smc5/6 functionality in rDNA stability, the lifespan of *smc6*-9 and *nse3*-1 mutants was characterized together with *fob1*Δ. Consistent with previous reports, the replicative lifespan of *fob*1Δ was extended compared to wild type^50, 52^. The reduced lifespan of *smc6*-9 was completely reversed by deletion of *FOB1*, as *smc6*-9 *fob*1Δ double mutants lived as long as *fob*1Δ (Fig. 5c). This is notable as deleting *FOB1* in *sir2*Δ mutants restored lifespan, but did not extend lifespan beyond wild type^51^. In stark contrast, the shortened lifespan of *nse3*-1, which was also accompanied by loss of silencing at NTS1 and NTS2 and destabilization of canonical silencing factor at NTS1, did not change in combination *fob*1Δ (Fig. 4c, Supplementary Fig. 4b). These data underscore the importance of Smc5/6 in rDNA stability independently of events at the RFB.

ERC formation, arising from recombination intermediates at Fob1-bound RFBs, is one measure of rDNA stability^29^. ERC levels increased in both *smc6*-9 and *nse3*-1 mutant cells (Fig. 5d). Previous work demonstrated that the production of ERCs can be reduced upon deletion of *FOB1* because forks no longer stall at the RFB^19, 50^. Consistent with this and the lifespan analysis, ERC formation in *smc6*-9 was dependent on *FOB1*, however ERCs persisted in *nse3*-1 *fob1*Δ double mutants (Fig. 5d). These data indicate that *fob1*Δ is able to rescue Smc5/6 HR-defects, but not Smc5/6 binding defects that contribute to replicative aging and ERC accumulation. Thus, the importance of the complex in rDNA stability is related to, but not completely dependent on, Fob1. Fob1-independent functions are perhaps linked to Smc5/6 at NTS2 as the level of Smc5^Myc^ recovered at NTS1 was reduced in cells where *FOB1* was deleted, however levels at NTS2 in *fob1*Δ were similar to wild type (Fig. 5e). Fob1 association at the rDNA was not altered in cells carrying either *nse3*-1 or *smc6*-9 (Supplementary Fig. 4a). Taken together, these data support a model for Smc5/6 in rDNA stability through binding NTS1 and NTS2, and that its role in transcriptional silencing and rDNA repeat compaction involve Smc5/6 binding at NTS2 independently of Fob1 (Fig. 6a).

**Fig. 6.**
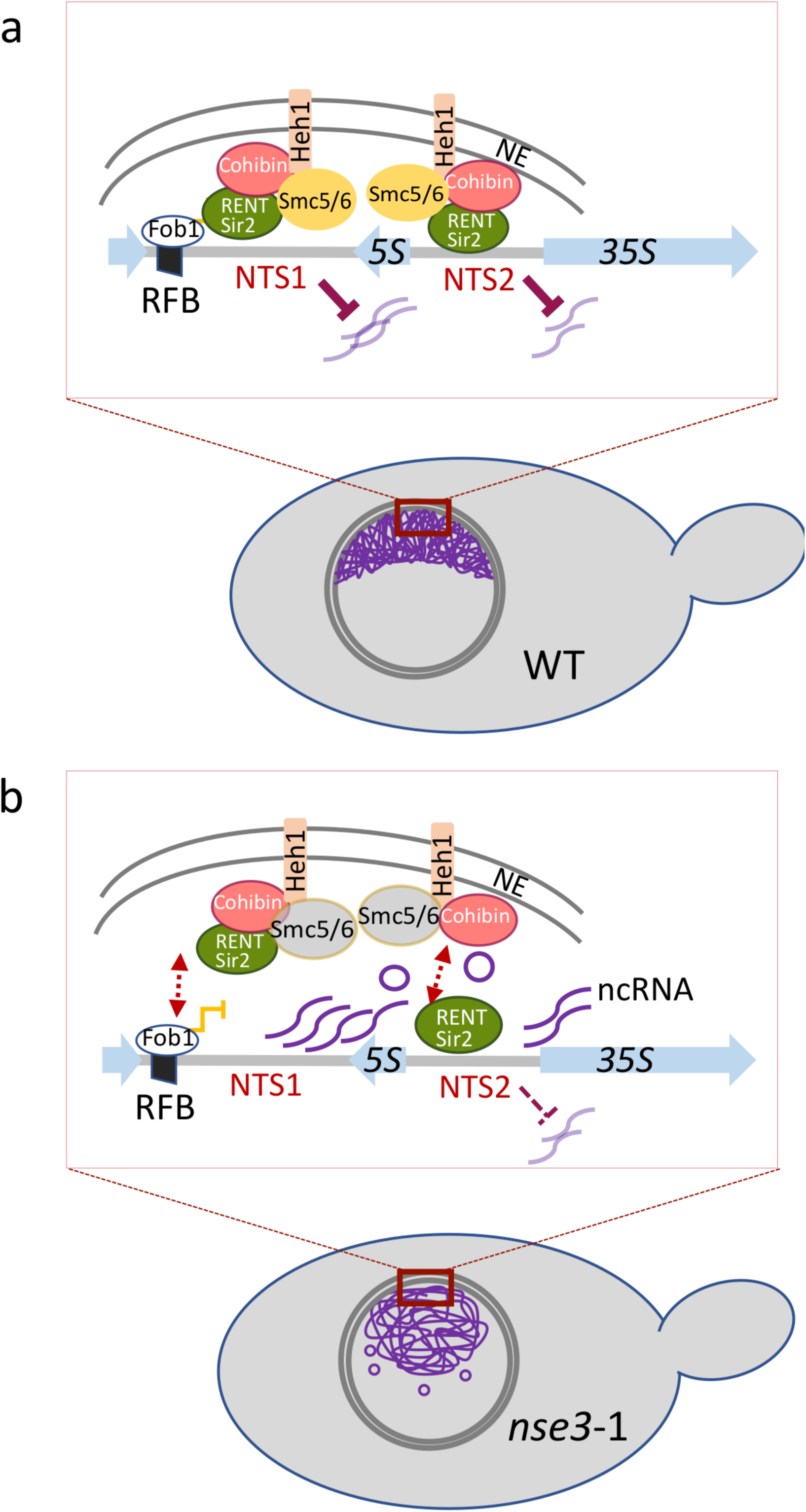
Schematic model representing interactions involved Smc5/6 complex to modulate silencing and morphology of the rDNA. In a WT cells, where Smc5/6 binds to the rDNA array, rDNA morphology is compact. Smc5/6 physically interacts with chromatin and canonical rDNA factors, Sir2, Cohibin and Heh1 whereby maintains silencing at NTS1 and NTS2. **b** In *nse3*-1 mutant, Smc5/6 fails to bind rDNA repeats, yet phisically interacts with Sir2, Cohibin and Heh1. Loss of the Smc5/6 complex results in defective silencing at both NTS1 and NTS2, accumulation of ERC molecules and increased nucleolar volume.

## DISCUSSION

The rDNA array in the yeast genome is unstable and we show that Smc5/6 is important for rDNA stability and maintaining replicative life span through two inter-related mechanisms involving: 1. transcriptional silencing of ncRNA at NTS1 and NTS2, and 2. tethering the rDNA repeats at the periphery. Moreover, its roles in silencing and nucleolar compartmentalization were uncoupled from its role in HR processing in experiments with separation-of-function mutants, *smc6*-9, which binds the rDNA but is HR deficient, and *nse3*-1, which does not associate with rDNA repeats.

The rDNA array is not static, but remains poised for repeat copy number contraction or expansion through a mechanism dependent on Fob1. Fob1 binding to the RFB is essential for programmed fork pausing and recruiting Sir2 as part of the RENT complex to silence chromatin in NTS1. The current study together with work by others demonstrate that Smc5/6 is involved in both of these processes. We show here that silencing at NTS1 is compromised in both *smc6*-9 and *nse3*-1 mutants and work from multiple labs have shown the importance of Smc5/6 in the resolution of HR structures that arise during programmed fork pausing^37, 38, 54^. Our data support previous work showing that Fob1-dependent fork pausing and transcriptional silencing at NTS1 are separately regulated^48^. Silencing defects in *smc6*-9 mutants increase in *smc6*-9 *fob1*Δ double mutant cells at NTS1. However, in this HR-deficient mutant, rDNA stability increased upon *FOB1* deletion, as the level of ERC accumulation decreased.

Smc5/6 interacts physically and genetically with Sir2 and Cohibin (Lrs4/Csm1), which are factors known to bind the NTS regions and silence chromatin. We show that like Sir2 and Cohibin, Smc5/6 also interacts with Heh1, tethering the rDNA repeats at the nuclear periphery^47, 51^. The binding of these canonical silencing complexes and Heh1 to the rDNA was markedly compromised in *nse3*-1 mutants. All of these complexes are known to bind other genomic loci including telomeres, centromeres and the mating type loci. As the association of Smc5/6 with these factors was unaltered in co-IPs in *nse3*-1 mutants, *in vivo* associations elsewhere might still persists^7, 24, 25, 34–36, 44, 55–58^.

Lifespan and rDNA instability are linked. One measure of rDNA stability is the formation of ERCs, which accumulated in both Smc5/6 mutants with reduced lifespans and was shorter in *nse3*-1 than in *smc6*-9 mutants. The deletion of *FOB1* reversed this in *smc6*-9, but *fob1*Δ did not impact the short lifespan of *nse3*-1. This observation together with ChIP of the canonical silencing factors and silencing data support a model whereby silencing in NTS2 correlates with lifespan independently of events at NTS1. First, *fob1*Δ cells with an increased lifespan show defects in silencing at NTS1, but not NTS2. Second, like *nse3*-1 and consistent with previous work, silencing at NTS2 is also disrupted in *sir2*Δ and to a lesser extent in *lrs4*Δ mutants with shortened lifespans^3–5, 7, 24, 25^. The deletion of *FOB1* in these mutants does not extend lifespan past wild type levels like it did in *smc6*-9 *fob1*Δ where NTS2 remained silenced. In *sir2*Δ mutants this discrepancy might be related to a loss of NTS2-bound Smc5/6, which was not disrupted in *smc6*-9 mutant cells. Importantly however, the binding of Sir2 and Cohibin at NTS1 are not limiting factors for lifespan extension^7, 48^ as these factors were equally down in *smc6*-9 *fob1*Δ, with an extended lifespan, and *nse3*-1 *fob1*Δ mutants, with a very short lifespan (Fig. 4c, Supplementary Fig. 5a). RNA polymerase I is essential for Sir2 binding to NTS2 and rDNA silencing^14, 59^. Therefore, investigating the interplay between RNA Pol I, Sir2 and Smc5/6 in the future will address further the importance of NTS2-based silencing on rDNA organization and lifespan.

Abnormal nucleolar morphology alteration has been demonstrated in mutants associated with rDNA instability, premature aging and in naturally aged cells^60–62^. The morphology of the nucleolus increased in *nse3*-1 mutant cells, however nucleolar size in *heh1*Δ remained similar to wild type. These data suggest that nucleolar volume increase correlates more with silencing rather than tethering as nucleolar size also increased in *sir2*Δ and *lrs4*Δ mutants with silencing defects, albeit to a lesser extent than in *nse3*-1. Moreover, our data support a model whereby morphology correlates more specifically with silencing defects at NTS2, which is high in *nse3*-1 and *nse3*-1 *fob1*Δ, but not in *smc6*-9 *fob1*Δ double mutants with compact nucleolar morphology. Even though nucleolar size in *heh1*Δ is similar to wild type, previous work shows that cells carrying the *HEH1* deletion have a shorter lifespan indicating that NTS silencing alone is insufficient for sustaining lifespan^24, 47^. Our data suggest Heh1-mediated tethering becomes important for rDNA stability when forks pause at the RFB. First, Heh1 association with the NTS regions is reduced in *fob1*Δ mutants with extended lifespan when forks do not pause (Supplementary Fig. 4c). Second, deletion of *FOB1* in *heh1*Δ mutants extended lifespan like it did when *fob1*Δ was combined with the HR-deficient *smc6*-9 allele. These findings are in agreement with a reduction in nucleolar size that accompanies lifespan extension after ectopic expression of a ‘rejuvenation factor’ in mitotically dividing aged yeast cells^45^.

The short lifespan of *nse3*-1 mutants correlated with loss of Smc5/6 binding in the rDNA and loss of transcriptional silencing at NTS2. Moreover, the nucleolar size increase and ERC accumulation that develops when Smc5/6 fails to localize to rDNA were not reversed by deletion of *FOB1*. We demonstrated that the physical association of Smc5/6 in the rDNA is important for silencing and nucleolar compaction, and that these functions correlate with rDNA stability and replicative lifespan. In all, our work support Smc5/6 as a structural maintenance of chromosome complex with involvement in mechanisms distinct of HR-processing at paused forks.

## Supporting information

Moradi-Fard Supp

## SUPPLEMENTAL INFORMATION

Supplemental information includes 4 figures and 3 tables and can be found with this article. Yeast strains used in this study are listed in Table S1. Details of plasmids and primers used in this study are specified in Table S2 and S3.

## ACKNOWLEDGEMENTS

We thank Dr. Karim Mekhail for reagents and helpful discussions. This work was supported by operating grants from CIHR MOP-82736; MOP-137062 and NSERC 418122 awarded to J.A.C. and CIHR and NSERC funding to T.A.A.H.

## AUTHORS CONTRIBUTIONS

S.M-F., A.M., M.C., and T.A.A.H. performed experiments and analyzed the data. J.A.C, S.M-F, and A.M. designed experiments and wrote the manuscript.

## METHODS

### EXPERIMENTAL MODEL AND SUBJECT DETAILS

All the yeast strains used in this study are listed in Table S1 and were obtained by crosses. The strains were grown on various media for the experiments, and are described below. For all experiments filter sterilized YPAD (1% yeast extract, 2% bactopeptone, 0.0025% adenine, 2% glucose and 2% agar) media were used. For yeast 2-hybrid assays, standard amino acid drop out media lacking histidine, tryptophan and uracil were used and 2% raffinose was added as the carbon source for the cells. In all experiments, exponentially growing cells were incubated at 30°C for 2hrs before harvesting, unless indicated otherwise.

#### Chromatin Immunoprecipitation

ChIP experiments performed as described previously 63. Cells were grown over night at 25°C, then diluted to 1×10^7^ cells/ml in liquid YPAD and incubated at 30°C for 2 hours before crosslinking with 1% formaldehyde (Sigma) for 15 minutes followed by quenching with 125 mM glycine for 5 minutes at room temperature. Fixed cells were washed 3 times with cold PBST (phosphate buffered saline with Tween 20) and froze over night at −80°C. Cells were lysed in lysis buffer (50 mm HEPES, 140 mm NaCl, 1 mm EDTA, 1% Triton X-100, 1 mM PMSF and protease inhibitor pellet), the clarified by spinning at 13200 rpm for 15 min (at 4°C). Pellets were sonicated for 12 x 15 seconds at amplitude of 50% with 45 seconds shut off intervals and immunoprecipitated using corresponding antibodies. Precipitates were washed once with lysis buffer and twice with wash buffer (100 mM Tris (pH 8), 0.5% Nonidet P-40, 1 mM EDTA, 500 mM NaCl, 250 mM LiCl, 1 mM PMSF and protease inhibitor pellet (Roche)) at 4°C, each for 5 minutes shaking at 1400 rpm. Real-time qPCR reactions were carried on using power up SYBR green master mix on a QuantStudio™ 6 Flex Real-Time PCR System (Applied Biosystems, Life Technologies Inc.). Ct (cycle threshold) values of Ab-coupled beads and uncoupled beads used to calculate fold enrichment of protein on rDNA regions relative to an unrelated genomic locus ZN (for ChIP experiments), or *ACT1* (for expression at rDNA).

#### Co-immunoprecipitation

Strains were grown overnight at 25°C and then diluted and grown to the log phase by incubating for 2 hours at 30°C in YPAD media. Cells were lysed with zirconia beads in lysis buffer (50 mm HEPES, 140 mm NaCl, 1 mm EDTA, 1% Triton X-100, 1mM PMSF and protease inhibitor pellet). Cell lysates were incubated with antibody-coupled Dynabeads for 2 hours at 4°C. Immunoprecipitates were washed end over end once with lysis buffer and twice with wash buffer (100 mM Tris (pH 8), 0.5% Nonidet P-40, 1 mM EDTA, 250 mM LiCl, 1 mM PMSF and protease inhibitor pellet), each for 5 minutes. Beads were resuspended in SDS loading buffer and subjected to SDS gel electrophoresis followed by western blotting using appropriate antibodies listed in the resource table.

#### qPCR based Gene Expression Analyses

Cells were grown over night at 25°C, then diluted to 5×10^6^ cells/ml in liquid YPAD and incubated at 30°C for 2 hours before fixing the cells with 1% Sodium azide. Fixed cells were washed with cold PBS (phosphate buffered saline; 1.37M NaCl, 27mM KCl, 100mM Na_2_HPO_4_, 18mM KH_2_PO_4_) and snap frozen in liquid nitrogen. Next day, cells were lysed using RNeasy kit reagents and isolated RNA was subjected to reverse transcription. Complementary DNA (cDNA) was amplified and quantified using the SYBR Green qPCR method. Primers are listed in Table S2. Expression values represent real time qPCR values relative to *ACT1* and normalization to WT samples.

#### PFGE

Saturated overnight culture cells were killed in 0.1% Sodium azide and washed with cold TE^50^ (10 mM Tris-HCl, pH 7.0, 50 mM EDTA, pH 8.0). To avoid mechanical shearing of genomic DNA, cells were solidified in 1% low melting-point CHEF-quality agarose in plug moulds (5×10^7^ cells/plug) at 4°C. Plugs were incubated overnight in 0.1 M sodium phosphate pH 7.0, 0.2 M EDTA, 40 mM DTT, 0.4 mg/ml Zymolyase 20T at 37°C, washed few times with TE50 and incubated in 0.5 M EDTA, 10 mM Tris–HCl pH 7.5, 1% *N*-lauroyl sarcosine, 2 mg/ml proteinase K for 48 hours at 37°C. Plugs were then washed with cold TE50 and stored at 4°C until subjected to electrophoresis. Chromosomes were separated on a CHEF-DRII instrument (Bio-Rad) for 68 hrs at 3.0 V/cm, 300–900 s, 14°C on a 0.8% CHEF agarose gel in 0.5% TBE. EtBr-stained gels were destained and then subjected to standard southern blotting as previously described (Moradi-Fard et al, 2016). Briefly, gels were treated with 0.25 N HCl for 20 min then in 0.5 M NaOH, 3 M NaCl for 30 min for in-gel depurinating and denaturing of genomic DNA respectively. Denatured DNA were transferred to Amersham Hybond-XL membrane overnight. Membranes were then crosslinked by UV Stratalinker 1800 (120 mJoules) and hybridized with radiolabeled rDNA specific probe^45^. Rediprime II DNA Labeling System used to radiolabel rDNA probe.

#### Quantification of ERC molecules

Genomic DNA were prepared using standard protocol. ∼2µg of DNA used to run in 0.8% agarose gel; 0.5x TBE. DNA fragments were separated for ∼24h at 40V, 4°C. Gels were then subjected to standard southern blotting and probed with rDNA-specific probe as described in PFGE section. ERC molecules were measured and represented after normalizing to genomic rDNA band using the ImageJ software.

#### Microscopy

Cells were grown overnight at 25°C and diluted to 5×10^6^ cells/ml and grown at 30°C to reach a concentration of 1×10^7^ cells/ml. Cells were washed twice with SK buffer (0.05M Kb_2_PO_4_, 0.05M K_2_HPO_4_, 1.2M Sorbitol). And mounted on slide for imaging. 15 Z-stack images were obtained with 0.3µm increments along the z-plane to cover a total range of cells nuclei at 60x magnification and 1.5µm/pixel zoom factor.

Three dimensional (X, Y, Z) stacks of yeast cells carrying Nop1-CFP and/or Nup49-GFP were acquired using the “Nikon Ti Eclipse Widefield” microscope provided by Live Cell Imaging facility at University of Calgary; ∼200ms and 400ms exposure times used for GFP and CFP channels respectively. The acquired 3D stacks were first deconvolved using Huygens software. 3D segmentation was done by thresholding (using the auto thresholding range recommended) in the ImageJ software using 3D manager plugin. The volume measurements were acquired in pixel and presented as relative to the obtained average volume (in pixel) for WT cells.

#### Yeast 2-hybrid

Various plasmids (Table S3) were constructed containing the gene encoding the proteins – Smc5, Nse1, Mms21, Nse3, Nse4, Nse6, Csm1, Lrs4 and Heh1-using the primers listed in Table S2. The plasmids J 965 and J 1493 and the inserts were treated with corresponding enzymes and ligated using T4 DNA ligase. The plasmids were sequence verified. Reporter (J 359), bait (J 965) and prey (J 1493) plasmids, containing the gene encoding the desired protein, were transformed into JC 1280. Cells were grown overnight in media lacking uracil, histidine and tryptophan with 2% raffinose. Next day, cells were transferred into media lacking uracil, histidine and tryptophan with either 2% glucose or 2% galactose and grown for 6 hrs at 30°C. Cell pellets were resuspended and then permeabilized using 0.1% SDS followed by ONPG addition. β-galactosidase activity was estimated by measuring the OD at 420nm, relative β-galactosidase units were determined by normalizing to total cell density at OD600.

#### Western Blot

Cells were lysed by re-suspending them in lysis buffer (with PMSF and protease inhibitor cocktail tablets) followed by bead beating with zirconia beads. The protein concentration of the whole cell extract was determined using the NanoDrop (Thermo Scientific). Equal amounts of whole cell extract were added to SDS PAGE gel wells. Standard SDS PAGE protocol were performed. Proteins were then transferred to nitrocellulose membrane and detected using corresponding antibodies listed in the resource table.

#### Quantification and Statistical Analysis

Data in bar graphs represent the average of at least 3 biological replicates. Error bars represent the standard error of mean (SEM). Significance (p value) was determined using 1-tailed, unpaired Student’s t test - p < 0.05^∗^; p < 0.01^∗∗^; p < 0.001^∗∗∗^. Statistical analyses were performed in Prism7 (GraphPad).

#### Data and Code Availability

This study did not generate/analyze any code. Original data supporting the figures in the paper is available from the corresponding author on request.

