## Supplementary material for "A Fob1 independent role for the Smc5/6 complex in the rDNA that modulates lifespan": Moradi-Fard Supp

Table 1: Strains used in this study

Table 2: Oligos used in this study

Table 3: Plasmids used in this study

Figures S1- S4

Table S1: Strains used in this study

| ***Strain*** | ***Genotype*** | ***Source*** |
|---|---|---|
| JC470 | *MAT***a** *ade2-1 trp1-1 his3-11 his3-15 ura3-1 leu2-3 leu2-112 Rad5+ (W303)* | R. Rothstein |
| JC471 | *MAT*α *ade2-1 trp1-1 his3-11 his3-15 ura3-1 leu2-3 leu2-112 Rad5+ (W303)* | R. Rothstein |
| JC1280 | *MAT***a** *his3 trp1 ura3-52 leu2::proLEU2-lexAop6* | Golemis, *et al*, 1996 |
| JC1358 | JC471 with *smc6*-9::*KanMX6* | C. Boone Lab |
| JC1594 | JC470 with Smc6-6HIS-3FLAG::*KanMX4* | Bustard *et al.,* 2012 |
| JC1595 | JC471 with Smc6-6HIS-3FLAG::*KanMX4* | this study |
| JC1725 | JC471 with Smc5-13Myc::*KanMX6*, *smc6*-9::*KanMX6* | this study |
| JC3032 | JC471 with *nse3*-1::*HYG* | Moradi-Fard *et al*., 2016 |
| JC3078 | JC1595 with *nse3*-1::*URA3* | this study |
| JC3467 | JC470 with Smc5-13Myc::*KanMX6* | this study |
| JC3483 | JC3467 with *nse3*-1::*URA3* | this study |
| JC3787 | JC470 with *sir2*::*TRP1*, *nse3*-1::*HYG* | this study |
| JC3790 | JC471 with *lrs4*::*KanMX6* | this study |
| JC3791 | JC470 with *lrs4*::*KanMX6* | K. Mekhail lab |
| JC3796 | JC3791 with *nse3*-1::*HYG* | this study |
| JC4022 | JC470 with Heh1-13Myc::*KanMX6* | this study |
| JC4107 | JC470 with Heh1-TAP::*TRP1,* TELVR::*ADE2* | K. Mekhail lab |
| JC4195 | JC470 with Csm1-TAP::*TRP1* | K. Mekhail lab |
| JC4196 | JC471 with Csm1-TAP::*TRP1* | K. Mekhail lab |
| JC4205 | JC1595 with *heh1*::*HYG* | this study |
| JC4224 | JC471 with Heh1-13Myc::*KanMX6* | this study |
| JC4228 | JC4022 with *nse3*-1::*HYG* | this study |
| JC4233 | JC471 with Csm1-TAP::*TRP1* | K. Mekhail lab |
| JC4243 | JC1595 with *csm1*::*KanMX6* | this study |
| JC4251 | JC4233 with *nse3*-1::*HYG* | this study |
| JC4399 | JC471 with *fob1::HIS3* | this study |
| JC4595 | JC3032 with *fob1::HIS3* | this study |
| JC4598 | JC1594 with Csm1-TAP::*TRP1* | this study |
| JC4648 | JC471 with *sir2*::*TRP1* | this study |
| JC4676 | JC471 with Nup49-*GFP*, Nop1-CFP::*URA3* | this study |
| JC4699 | JC1595 with *sir2*::*TRP1* | this study |
| JC4712 | JC4598 with *nse3*-1::*HYG* | this study |
| JC4729 | JC470 with Nup49-*GFP*, Nop1-CFP::*URA3*, *nse3*-1::*HYG* | this study |
| JC4731 | JC4676 with *lrs4*::*KanMX6* | this study |
| JC4733 | JC4676 with *sir2*::*TRP1* | this study |
| JC4735 | JC4676 with *heh1*::*HYG* | this study |
| JC4773 | JC4774 with *nse3*-1::*HYG* | this study |
| JC4774 | JC4224 with Csm1-TAP::*TRP1* | this study |
| JC4811 | JC1595 with Heh1-TAP::*TRP1* | this study |
| JC4813 | JC4811 with *nse3*-1::*URA3* | this study |
| JC4824 | JC1358 with *fob1::LEU2* | this study |
| JC4825 | JC471 with *fob1::LEU2* | this study |
| JC4929 | JC4824 *with* Csm1-TAP::*TRP1* | this study |
| JC4932 | JC4676 with *smc6*-9::*KanMX6* | this study |
| JC4937 | JC4233 with *fob1:: LEU2* | this study |
| JC4938 | JC4233 with *smc6*-9::*KanMX6* | this study |

| | | |
|---|---|---|
| JC4940 | JC4022 with *fob1::LEU2* | this study |
| JC4942 | JC4022 with *smc6*-9::*KanMX6* | this study |
| JC4943 | JC4022 with *smc6*-9::*KanMX6, fob1::LEU2* | this study |
| JC4973 | JC4233 with *fob1::LEU2*, *nse3*-1::*HYG* | this study |
| JC4975 | JC4022 with *fob1::LEU2*, *nse3*-1::*HYG* | this study |
| JC4976 | JC1358 with *lrs4*::*KanMX6* | this study |
| JC4978 | JC1358 with *sir2*::*HIS3* | this study |
| JC4979 | JC3790 with *sir2*::*HIS3* | this study |
| JC4980 | JC4979 with *nse3*-1::*HYG* | this study |
| JC4985 | JC5016 with *fob1::LEU2* | this study |
| JC5007 | JC471 with Fob1-3HA:: *HIS3* | this study |
| JC5008 | JC5007 with *nse3*-1::*HYG* | this study |
| JC5010 | JC5007 with *smc6*-9::*KanMX6* | this study |
| JC5014 | JC5016 with *smc6*-9::*KanMX6* | this study |
| JC5015 | JC5016 with *nse3*-1::*HYG* | this study |
| JC5016 | JC471 with Nop1-CFP::*URA3* | this study |
| JC5017 | JC5016 with *sir2*::*TRP1* | this study |
| JC5018 | JC5016 with *heh1*::*HYG* | this study |
| JC5019 | JC5016 with *lrs4*::*KanMX6* | this study |
| JC5039 | JC3467 with *smc6*-9::*KanMX6* | this study |
| JC5040 | JC5039 with *fob1::LEU2* | this study |
| JC5041 | JC3467 with *fob1::LEU2* | this study |
| JC5044 | JC3483 with *fob1::LEU2* | this study |
| JC5110 | JC5015 with *fob1::HIS3* | this study |
| JC5113 | JC5014 with *fob1::HIS3* | this study |

Table S2: Oligos used in this study

| ***Primer name*** | ***Primer number*** | ***Sequence 5'> 3'*** |
|---|---|---|
| NTS1 | C1577 | AGGGCTTTCACAAAGCTTCC |
| | C1578 | TCCCCACTGTTCACTGTTCA |
| NTS2 | C1795 | CCACCACACTCCTACCAATAAC |
| | C1796 | AGGTAGTCAGATGAAAGATGAATAGAC |
| P1 | C1791 | CACACTATCATCCTCATCGTATATT |
| | C1792 | AGAGAGAAGTAGACTGAACAAGT |
| P2 | C1799 | ACGATGAGAGACTGTTCAGGTTAAA |
| | C1800 | GGGTTGATGCGTATTGAGAGATA |
| P3 | C1801 | CCAATTGTTCCTCGTTAAGGTATTT |
| | C1802 | ATTCAGGGAGGTAGTGACAATAAA |
| P4 | C1807 | GTTTGAGAATAGGTCAAGGTCATTTC |
| | C1808 | GTTTCCCTCAGGATAGCAGAAG |
| ZN | C1275 | GCACTTAATTGGCGTAAGCTG |
| | C1276 | TCGCAGGAGCATATTTCGTA |
| Act1 | C1561 | TGTCCTTGTACTCTTCCGGT |
| | C1562 | CCGGCCAAATCGATTCTCAA |
| Smc5 | C1483 | GATCCCATGGATGACCAGTCTAATAGATTTGGGCAGATATG |
| | C1484 | GATCCTCGAGTTAATCGAATGAGTAGTTAGAAGTTTCACCG |

| | | |
|---|---|---|
| Nse1 | C719 | CTAGGAATTCATGGAGGTACATGAAGAGC |
| | C720 | CTAGCTCGAGTTAAATAACGTATACGCCCTCTG |
| Nse3 | C723 | CTAGGAATTCATGAGTTCTATAGATAATGAC |
| | C724 | CTAGCTCGAGCTATATAGAATATGAATCGCC |
| Nse4 | C721 | CTAGGAATTCATGTCTAGTACAGTAATATC |
| | C722 | CTAGCTCGAGTAAGAATGGTGAAGTGATGTTG |
| Nse6 (pJ1493) | C609 | GATCGGATCCGTGTCACAAATGGGAAGCGTGAACTCATCACCG |
| | C610 | GATCCTCGAGCAGATCAATGTTCAGTCATCATGACTGTTACC TG |
| Nse6 (pJ965) | C892 | GATCGGATCCAAATGGGAAGCGTGAACTCATCACCG |
| | C610 | GATCCTCGAGCAGATCAATGTTCAGTCATCATGACTGTTACC TG |
| Csm1 | C1737 | GATC GAATTC ATGGATCCATTGACTGTATACAAAAACTCAGTGAAACA |
| | C1738 | GATC CTCGAG TTATGTAGCAGCTTACTCGGTTTCATCTTTTTTCTCTC |
| Lrs4 | C1735 | GATCGAATTCATGGAGCATGTAGATTCCGATTTTGCACCTATAAGGAG |
| | C1736 | GATC CTCGAG GATAGCTGTTACTCATACAAACTCGTCAACATTTAAAT |
| Heh1 | C1898 | GATCGCGGCCGCATGAATAGTGACTTGGAGTATTTAGAGGACGGTTTTGA |
| | C1915 | GATCCTCGAGTCATTTTGTGGGTTATATTTTGTTTTCAGCGGAATCCT |

Table S3: Plasmids Used in this study

| ***Plasmid number*** | ***Plasmid description*** | ***Source*** |
|---|---|---|
| J 965 | pGAL-lexA | S. Gasser lab |
| J 1493 | pJG4-6 | S. Gasser lab |
| J 359 | pSH18-34 lexAGal1-lacZ | |
| J 1805 | J 965 with Nse1 | |
| J 102 | J 965 with Mms21 | |
| J 138 | J 965 with Nse3 | |
| J 1804 | J 965 with Nse4 | |
| J 141 | J 965 with Nse6 | |
| J 1902 | J 965 with Smc5 | |
| J 1874 | J 965 with Csm1 | |
| J 1875 | J 965 with Lrs4 | |
| J 1900 | J 1493 with Nse1 | |
| J 050 | J 1493 with Mms21 | |
| J 1816 | J 1493 with Nse3 | |
| J 1901 | J 1493 with Nse4 | |
| J 063 | J 1493 with Nse6 | |
| J 048 | J 1493 with Smc5 | |
| J 1881 | J 1493 with Heh1 | |
| J 187 | pFN4 pNOP1-CFP | #1742 S. Gasser lab |
| J 1830 | pNOY373, containing 9.1 Kb rDNA repeats | (Unal *et al.,* 2011) |

a

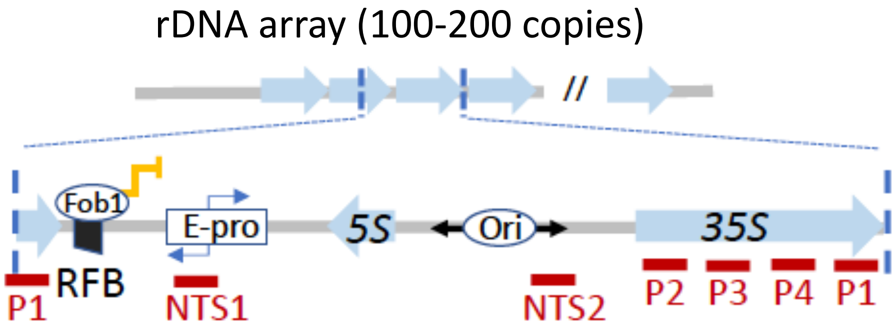


b

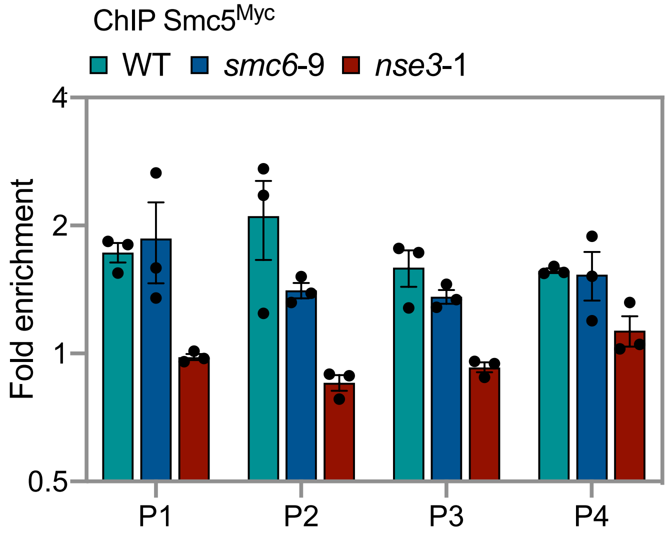


**Supplementary Fig. 1, Related to Figure 1.**
**a** Location of P1-P4 probes in the rDNA. **b** Enrichment of Smc5$^{Myc}$ at NTS1 and NTS2 by ChIP with α-Myc in at P1-P4 in WT (JC 3467), *nse3*-1 (JC 3483) and *smc6*-9 (JC 5039). **b** Fold enrichment is based on normalization to negative control region (ZN). Analysis was performed using at least three biological replicates. Statistical analysis is described in methods.

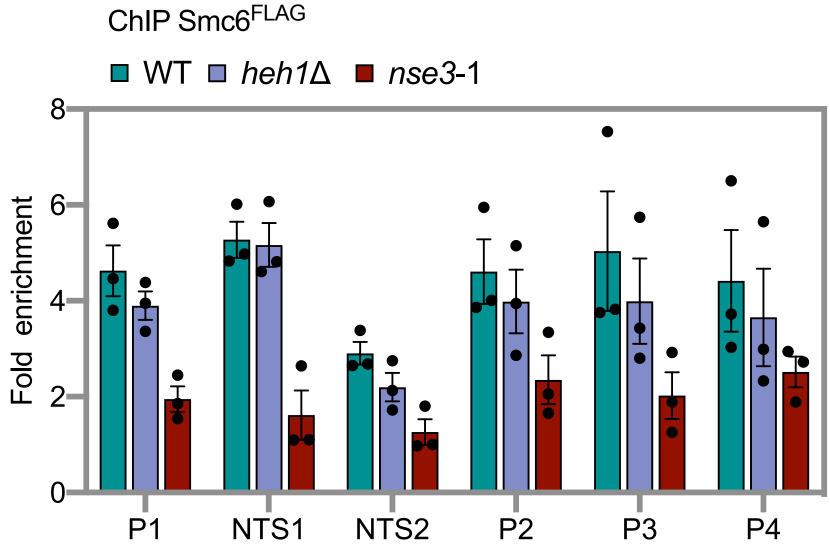


**Supplementary Fig. 2, Related to Figure 2.**
Enrichment of Smc6$^{FLAG}$ at NTS1 and NTS2 by ChIP with α-FLAG in WT (JC 1595), *heh1*Δ (JC 4205) and *nse3*-1 (JC 3078). Fold enrichment is based on normalization to negative control region (ZN). Analysis was performed using at least three biological replicates. Statistical analysis is described in methods.

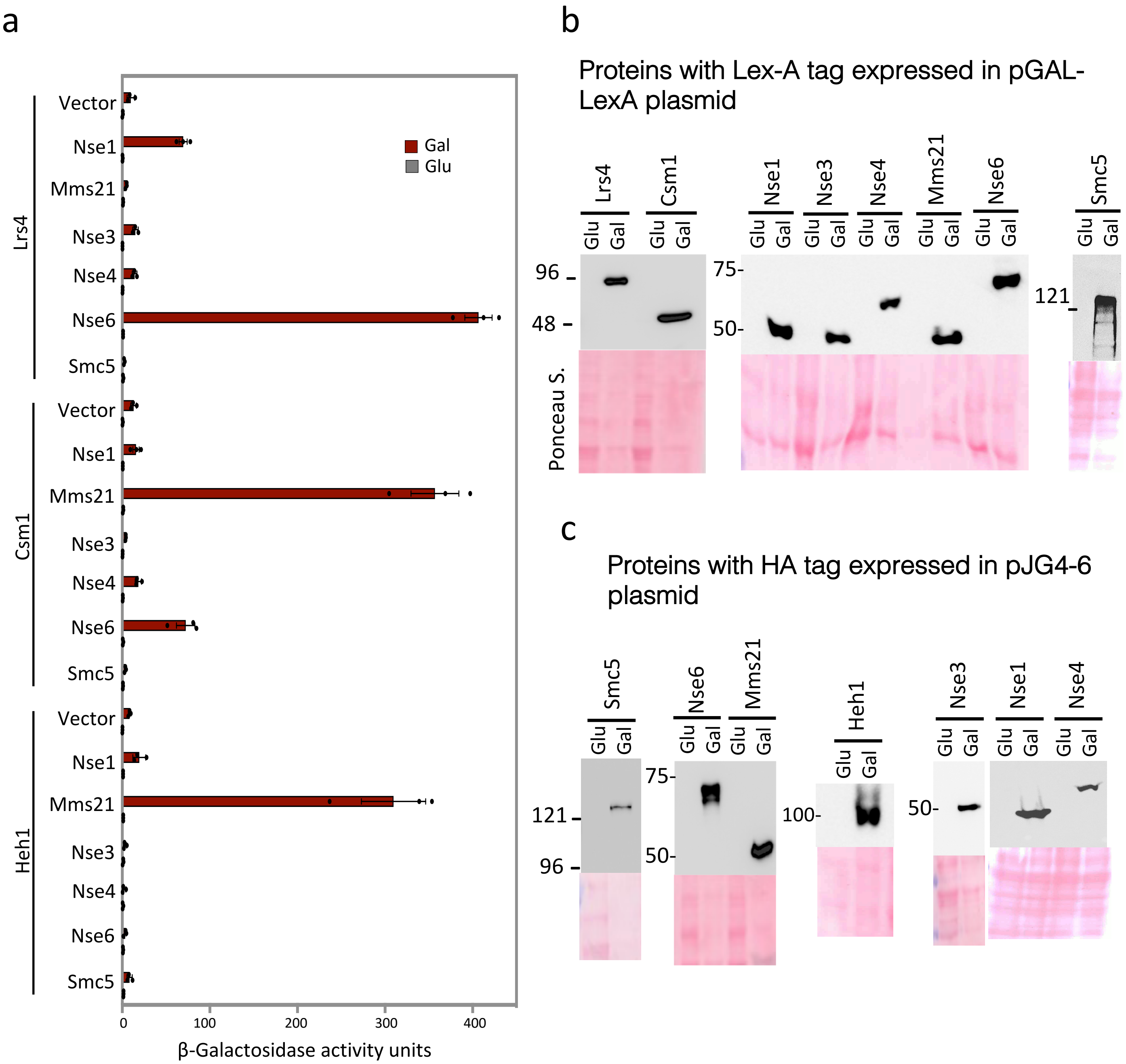


**Supplementary Fig. 3, Related to Figure3.**
**a** Yeast-two Hybrid analysis between Smc5/6 (Nse1, Mms21, Nse3, Nse4, Nse6 and Smc5), Cohibin (Lrs4 and Csm1) components and Heh1 using quantitative β-galactosidase activity assay. **b** Western blots with α-LexA and **c** α-HA shows the expression levels of proteins with their corresponding epitope tags from Yeast-two hybrid vectors after induction in Galactose-containing media. Glu: Glucose; Gal: Galactose. The expression vectors are listed in Table S3.

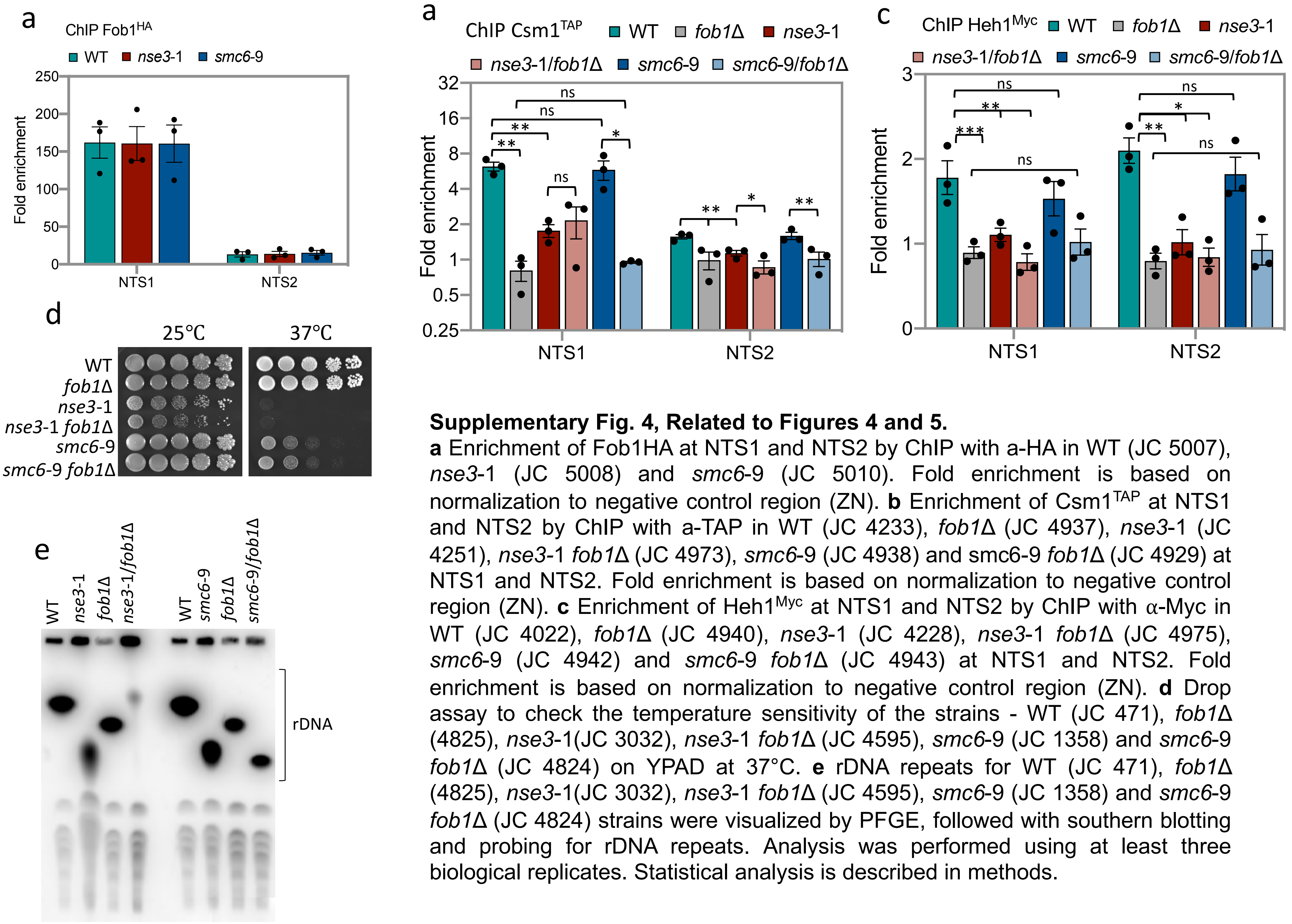


**Supplementary Fig. 4, Related to Figures 4 and 5.**

**a** Enrichment of Fob1HA at NTS1 and NTS2 by ChIP with a-HA in WT (JC 5007), *nse3*-1 (JC 5008) and *smc6*-9 (JC 5010). Fold enrichment is based on normalization to negative control region (ZN). **b** Enrichment of Csm1$^{TAP}$ at NTS1 and NTS2 by ChIP with a-TAP in WT (JC 4233), *fob1Δ* (JC 4937), *nse3*-1 (JC 4251), *nse3*-1 *fob1Δ* (JC 4973), *smc6*-9 (JC 4938) and smc6-9 *fob1Δ* (JC 4929) at NTS1 and NTS2. Fold enrichment is based on normalization to negative control region (ZN). **c** Enrichment of Heh1$^{Myc}$ at NTS1 and NTS2 by ChIP with α-Myc in WT (JC 4022), *fob1Δ* (JC 4940), *nse3*-1 (JC 4228), *nse3*-1 *fob1Δ* (JC 4975), *smc6*-9 (JC 4942) and *smc6*-9 *fob1Δ* (JC 4943) at NTS1 and NTS2. Fold enrichment is based on normalization to negative control region (ZN). **d** Drop assay to check the temperature sensitivity of the strains - WT (JC 471), *fob1Δ* (4825), *nse3*-1(JC 3032), *nse3*-1 *fob1Δ* (JC 4595), *smc6*-9 (JC 1358) and *smc6*-9 *fob1Δ* (JC 4824) on YPAD at 37°C. **e** rDNA repeats for WT (JC 471), *fob1Δ* (4825), *nse3*-1(JC 3032), *nse3*-1 *fob1Δ* (JC 4595), *smc6*-9 (JC 1358) and *smc6*-9 *fob1Δ* (JC 4824) strains were visualized by PFGE, followed with southern blotting and probing for rDNA repeats. Analysis was performed using at least three biological replicates. Statistical analysis is described in methods.
